# Redundancy between spectral and higher-order texture statistics for natural image segmentation

**DOI:** 10.1101/2021.04.26.441524

**Authors:** Daniel Herrera-Esposito, Leonel Gómez-Sena, Ruben Coen-Cagli

**Affiliations:** Laboratorio de Neurociencias, Facultad de Ciencias, Universidad de la República, Montevideo, Uruguay; Dept. of Systems and Computational Biology and Dominick P. Purpura Dept. of Neuroscience, Albert Einstein College of Medicine, Bronx, NY, USA

## Abstract

Visual texture, defined by local image statistics, provides important information to the human visual system for perceptual segmentation. Second-order or spectral statistics (equivalent to the Fourier power spectrum) are a well-studied segmentation cue. However, the role of higher-order statistics (HOS) in segmentation remains unclear, particularly for natural images. Recent experiments indicate that, in peripheral vision, the HOS of the widely adopted Portilla-Simoncelli texture model are a weak segmentation cue compared to spectral statistics, despite the fact that both are necessary to explain other perceptual phenomena and to support high-quality texture synthesis. Here we test whether this discrepancy reflects a property of natural image statistics. First, we observe that differences in spectral statistics across segments of natural images are redundant with differences in HOS. Second, using linear and nonlinear classifiers, we show that each set of statistics individually affords high performance in natural scenes and texture segmentation tasks, but combining spectral statistics and HOS produces relatively small improvements. Third, we find that HOS improve segmentation for a subset of images, although these images are difficult to identify. We also find that different subsets of HOS improve segmentation to a different extent, in agreement with previous physiological and perceptual work. These results show that the HOS add modestly to spectral statistics for natural image segmentation. We speculate that tuning to natural image statistics under resource constraints could explain the weak contribution of HOS to perceptual segmentation in human peripheral vision.

## 1) Introduction

Scene segmentation is an essential function of visual processing. Grouping visual features together in a segment and separating different segments in a scene requires multiple processes and sources of information. These include gestalt principles such as proximity, similarity, common fate (Wagemans et al., 2012); geometrical cues such as symmetry and collinearity (Field et al., 1993; Geisler et al., 2001; Sigman et al., 2001); statistical cues related to texture information (Julesz, 1962; Landy & Bergen, 1991; Z. Li, 2002); binocular disparity cues (Bakin et al., 2000; Zhaoping et al., 2009); detection of edges and boundaries between regions (Ben-Shahar, 2006; Wolfson & Landy, 1998); and shape information derived from object recognition and semantic understanding of scenes (Neri, 2014, 2017), just to name a few.

Here we focus on visual texture, which can be defined by the local statistical properties of an image region (Julesz, 1962; Victor et al., 2017). This is a particularly important and well-studied substrate for image segmentation, as reflected in the vast perceptual (Julesz, 1962; Landy, 2013; Landy & Bergen, 1991; Victor, 1994; Victor et al., 2017) and physiological (Knierim & van Essen, 1992; V. A. Lamme, 1995; V. A. F. Lamme et al., 1999; Nothdurft et al., 2000; Roelfsema, 2006) literature and in successful models of human texture segmentation (Bergen & Landy, 1991; Bhatt et al., 2007; Z. Li, 2002; Malik & Perona, 1990; Victor et al., 2017). Notably, most of this work has been focused on studying second-order statistics (represented in the Fourier power spectrum, henceforth spectral statistics), despite abundant evidence that higher-order statistics (HOS) also strongly influence texture perception (Balas, 2006; Freeman et al., 2013; Freeman & Simoncelli, 2011; Hermundstad et al., 2014; Julesz et al., 1978; Portilla & Simoncelli, 2000; Tesileanu et al., 2020; Victor et al., 2013; Victor & Conte, 1996) and are essential to capture the appearance of natural textures (Balas, 2006; Portilla & Simoncelli, 2000). Studies of HOS cues for texture segmentation have used artificial textures, and relatively low-order statistics (Hermundstad et al., 2014; Julesz et al., 1978; Tesileanu et al., 2020; Tkacik et al., 2010; Victor et al., 2013; Zavitz & Baker, 2014). As a consequence, the relevance of HOS for texture-based segmentation remains uncertain, particularly in the context of natural vision.

Texture processing is especially prominent in peripheral vision, and the most influential theory of peripheral vision relies on summary statistics (SS) of textures (Balas et al., 2009; Freeman et al., 2013; Freeman & Simoncelli, 2011; Rosenholtz, 2016). One important instantiation of the SS theory relies on the statistics defined by the Portilla-Simoncelli (PS) algorithm for texture synthesis (Balas et al., 2009; Freeman & Simoncelli, 2011; Portilla & Simoncelli, 2000), which uses marginal pixel statistics, spectral statistics and a specific set of HOS (detailed below) to synthesize textures with naturalistic appearance. The PS instantiation of the SS model uses a filtering stage analogous to the primary visual cortex (V1) followed by a HOS encoding stage (**Figure 1b**), and captures many aspects of peripheral perception (Balas et al., 2009; Ehinger & Rosenholtz, 2016; Freeman et al., 2013; Freeman & Simoncelli, 2011; Rosenholtz, 2016; Rosenholtz et al., 2012) as well as the selectivity of neurons at higher stages of the visual cortex (Freeman et al., 2013; Okazawa et al., 2015, 2017; Ziemba et al., 2016).

**Figure 1.**
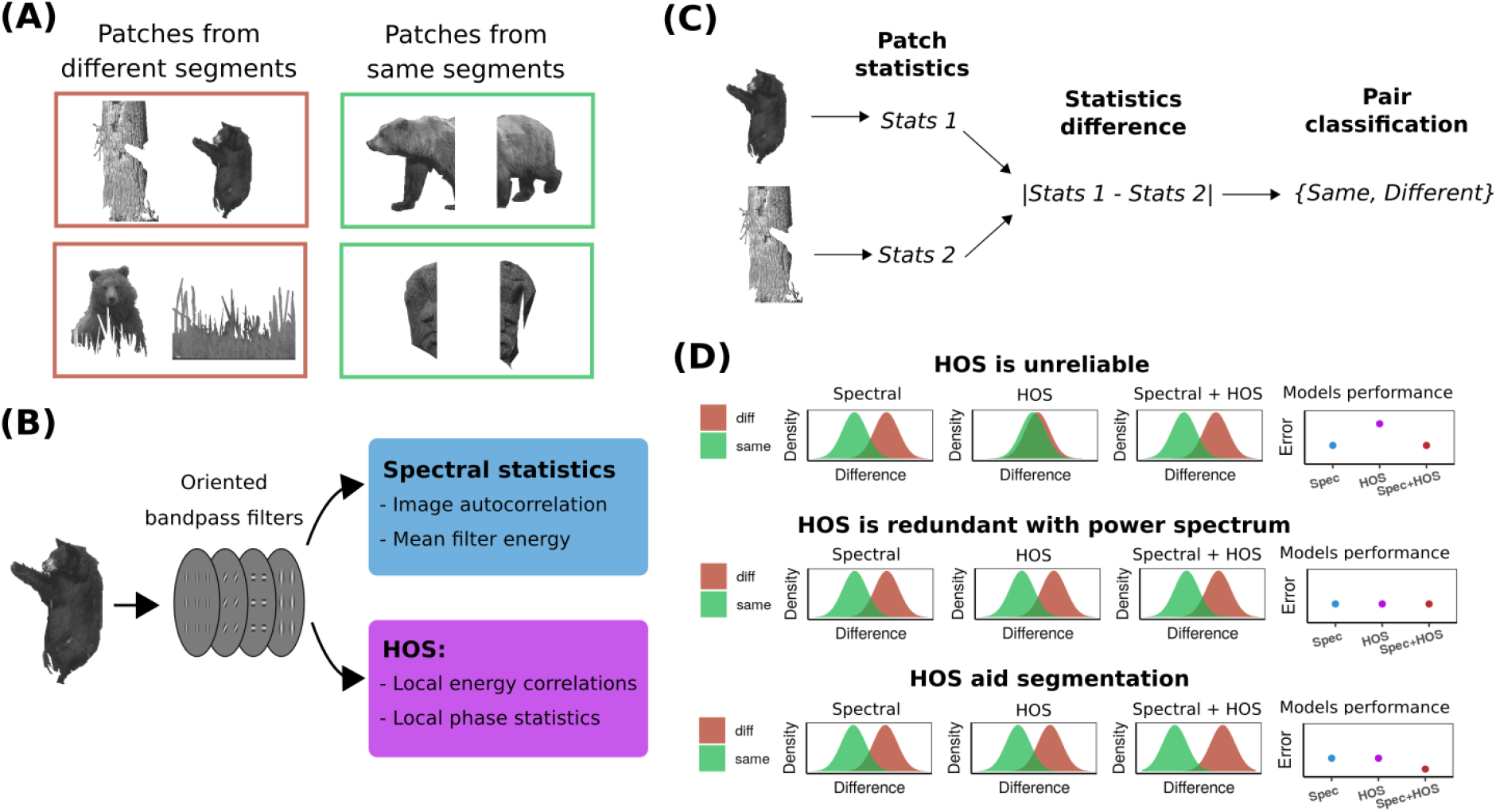
(A) Example pairs of patches taken from different (left) or from the same (right) image segment of a natural image. (B) Illustration of the computing of the different image statistics, with a first filtering stage and a second stage of computing image statistics. (C) Segmentation task, in which the statistics of two image patches are used to classify them as belonging to the same or to different segments. (D) Each row illustrates one possible scenario of spectral statistics and HOS contributions to image segmentation. Plots show the distribution of the difference between statistics across image patches from the same (green) and from different (brown) segments for different combinations of statistics (first three columns), and the corresponding performance of different segmentation models using these statistics (fourth column).

Recent work by us and others (Doerig et al., 2019; Herrera-Esposito et al., 2021; Herzog et al., 2015; Manassi et al., 2012, 2013; Wallis et al., 2019) suggests that perceptual segmentation is an important missing component from the SS model of the visual periphery. In particular, we (Herrera-Esposito et al., 2021) observed that segmentation cues improve performance in a naturalistic texture discrimination task, when target textures are surrounded by distractor textures. Remarkably, however, when we introduced a difference in HOS between target and distractor textures, that difference induced little segmentation, on average, if these regions shared the same spectral statistics (although there is some between texture variability in the estimated effect of HOS). This observation raises the following question: why are these HOS only a weak segmentation cue (relative to spectral statistics) to our peripheral visual systems?

Here we test whether this observation reflects a property of natural images’ statistics, which may be exploited by the human peripheral visual system through processes of statistical inference (Hindi Attar et al., 2007), or whether it reflects suboptimal processing. Under the first hypothesis, two scenarios are possible. First (**Figure 1D**, top), the HOS might be an unreliable cue for texture-based segmentation, because differences in local HOS between two regions do not reliably correspond to differences in the segments of the scene. In this case, the visual system would learn to weigh the spectral statistics information more heavily than the HOS information, similar to much previous research in visual (Adams & Mamassian, 2004; Jacobs, 1999; Knill & Saunders, 2003; Saarela & Landy, 2012), auditory (Cazettes et al., 2014; Pavão et al., 2020) and multisensory (Fetsch et al., 2012; Gu et al., 2008) cue combination. A second possibility (**Figure 1D**, middle) is that the HOS may be a reliable cue for segmentation, but highly redundant with the spectral statistics. For example, if it seldom occurs that two different neighboring segments in a natural image have similar spectral statistics but different HOS that allow to segregate them, then these HOS would add little information to the process of texture-based segmentation of natural images. Then, using the spectral statistics but not HOS for peripheral segmentation, could be advantageous considering resource constraints (see Discussion). Lastly (**Figure 1D**, bottom), an alternative hypothesis is that both spectral statistics and HOS are informative about segmentation and independent of each other, in which case the smaller weight placed on HOS by peripheral segmentation processes would reflect a combination of inaccurate encoding of the HOS of PS and suboptimal readout of that information.

To test those possibilities, first we studied how spectral statistics and HOS change across natural textures and natural scenes segments. Next, we trained an observer model to solve a classification task using different combinations of spectral statistics and HOS, in which the goal is to determine whether two image patches belong to the same image segment or not (see **Figure 1**). We used both images of composite natural textures, where we defined the ground-truth segmentation, and images of natural scenes with segmentation maps drawn by humans (Martin et al., 2001). Our results provide the first quantification of the relative power of spectral statistics and HOS of the PS model for texture-based segmentation of natural images.

## 2) Methods

### 2.1) Image and segment selection

For the analysis of texture images we used 638 natural texture images obtained from the Brodatz (Brodatz, 1966), Salzburg Texture Image (*Salzburg Texture Image Database (STex)*, n.d.), and the Lazebnik et al. databases (Lazebnik et al., 2005). We converted the color images to grayscale with the *image* package for octave, by extracting the luminance channel of the YIQ color space. We then normalized the pixel values of each image to have a mean of 0.5 and a standard deviation of 0.2 on a scale between 0 and 1. Next, we cropped 4 non-overlapping square patches of 128 x 128 pixels from the vertices of the image, thus obtaining 4 sample patches per texture (a total of 2552 patches).

For the natural scene analysis we used the 500 natural scene images from the Berkeley segmentation database (BSD) (Martin et al., 2001), and their corresponding segmentation maps labeled by a human (we used the first map available for each image). We converted the color images to grayscale with the same procedure as for textures. The segments analyzed for each image were the central segment of the image (the one containing the central pixel) and all its neighbors. To avoid excessive noise in the computed statistics, we filtered out the images in which the central segment had less than 8192 pixels (equivalent to a 128 x 64 pixels). Furthermore, we also filtered out the neighboring segments with less than 4096 pixels (equivalent to a 64 x 64 pixel patch). After this selection procedure, 416 images and a total of 1696 segments were used.

### 2.2) Pairing image patches and texture segmentation task

Region-based segmentation consists in the process of determining whether two image regions belong to the same segment or not. We modeled this region-based segmentation task using texture as a substrate by employing a classification task on pairs of image patches.

We generated pairs of image patches that could either belong to the same segment or to different segments (**Figure 1**). For brevity, we refer to these pairs as “matched” and “unmatched” respectively. Then, we computed the statistics of the patches (see details below) and, depending on the analysis, we either computed the angle between the vectors of statistics of the two patches (used in **Figure 2**; see subsection 2.3 for details), or we computed the absolute difference between the patches for each of the PS statistics (used in all other figures and tables). The classification task consists in determining whether the two texture patches belong to the same segment or not.

**Figure 2.**
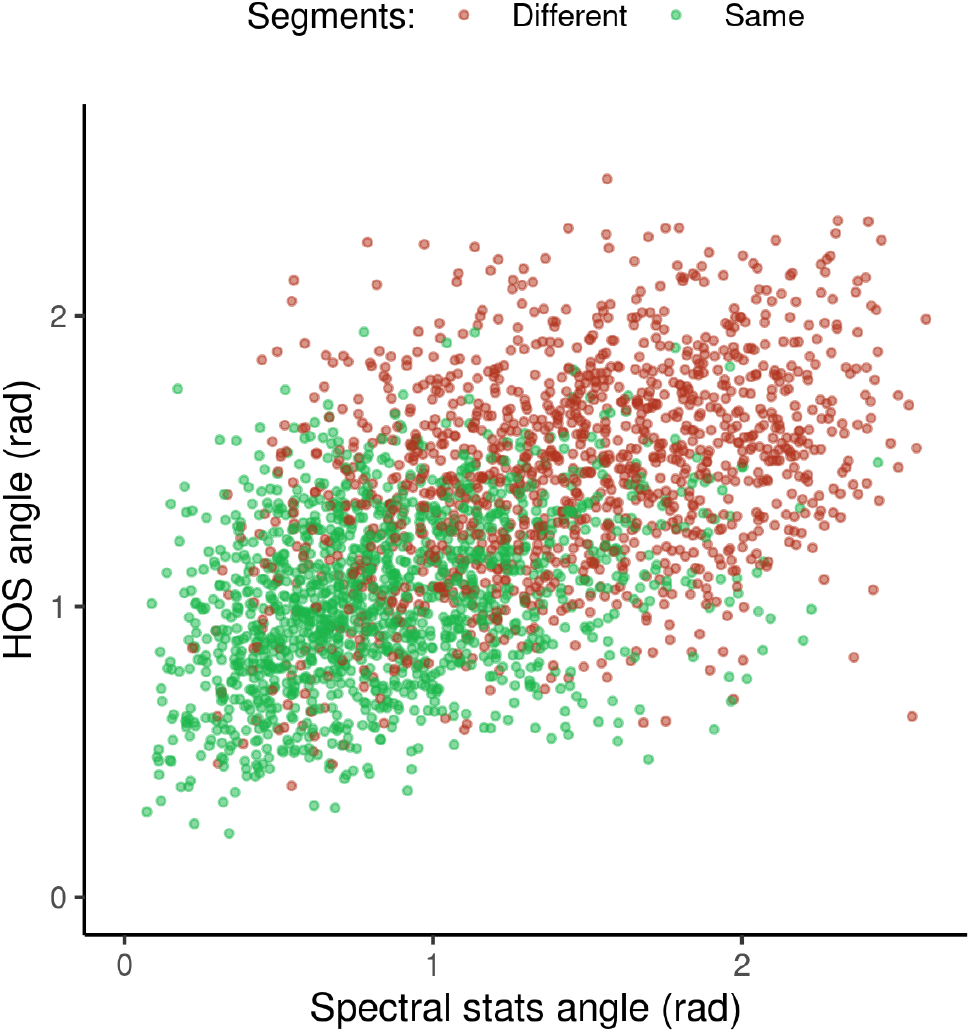
Relation between the difference in HOS and the difference in spectral statistics for pairs of image patches from the BSD database. The color of the dots indicates whether the pair of patches are extracted from the same image segment (green) or not (brown).

For the texture images we considered the whole image as one segment, and thus built the matched pairs by pairing two patches from the same texture, and the unmatched pairs by pairing two patches from different textures.

For the natural images in the BSD, the matched pairs were obtained by splitting the center patch, and the neighboring patches with more than 8192 pixels, vertically into two halves at the point that produced the most balanced pixel distribution, and pairing the two halves. The unmatched pairs were obtained by pairing the central patch of an image with a neighboring segment.

### 2.3) Computing texture statistics

For each image patch we computed the summary statistics of the PS model. These comprise a set of marginal pixel statistics, and a set of statistics over the filter outputs of the steerable pyramid (Portilla & Simoncelli, 2000). We used a bank of filters with 4 orientations and 4 scales, and a neighborhood of 7 pixels for computing spatial correlations. For the cropped texture patches we used the original PS code. For the patches of natural scenes we used a modified version of the Freeman metamer model (Freeman & Simoncelli, 2011). The original code first filters an image with the steerable pyramid, and then computes the weighted average of the pairwise products of filter outputs (equivalent to computing correlations), using a predetermined set of regular weighting windows. Our modification consisted in using an irregular weighting window, given by the segmentation map of the BSD image. In both textures and natural scenes we also modified the code to compute correlations where the original models computed covariances because we observed that correlations afforded better performance in the discrimination task.

We separated the statistics of the PS model into 3 groups, following previous work (Portilla & Simoncelli, 2000; Ziemba et al., 2016): pixel statistics, Fourier power spectrum (spectral statistics), and statistics of higher-order (HOS). The pixel statistics are marginal statistics over the pixel values, including mean, variance, and the skewness and kurtosis at different lowpass versions of the image. The spectral statistics are equivalent to pairwise pixel correlations, which are found in the PS model in the central autocorrelation matrices of the image subsampled at different scales, and in the mean modulus of activation of quadrature pairs of complex filters. The rest of the statistics in the PS model, which are not captured by the marginal pixel statistics or by pairwise pixel correlations, are referred to as HOS. These comprise correlations across space, scale, and orientation between the magnitude of complex bandpass quadrature filters (i.e. the energy of the filters), and local phase statistics (Portilla & Simoncelli, 2000). With the parameters we used for the PS model, we obtained 16 pixel statistics, 137 spectral statistics statistics, and 552 HOS.

### 2.4) Correlation between spectral statistics and HOS

To analyze the correlation between spectral statistics and HOS differences between image regions, we first z-scored each statistic across the BSD patches to have 0 mean and a standard deviation of 1. Then for each pair of patches we computed the angle between their vectors of spectral statistics and the angle between their vectors of HOS statistics (i.e. we computed the angles in the respective 137 and 552 dimensional spaces for the two kinds of statistics). Then we computed the Pearson correlation between these two.

### 2.5) Training the linear classifier models

All linear classification models using the absolute differences in statistics were trained by ridge regression, using the glmnet package v4.0-2 (Friedman et al., 2019) in R 3.6.3 (R Core Team, 2018). We used the default settings of the package in which the scaling parameter for the penalization is selected by 10-fold cross-validation on the training set. We used misclassification rate as the criterion for both selecting the penalization parameter and training the model. We also performed a weighting of the pairs of images in the training set so that the overall training weight was the same for the two classes. We also normalized each predictor to have unit variance and zero mean in the training set. We performed this procedure both for the models performing the segmentation task, as for the models performing the identification of pairs of patches with useful HOS.

### 2.6Training the segmentation models

For the image segmentation task, we trained a family of linear models to classify the pairs of patches using the absolute difference in each statistic between the patches. The subsets of the PS statistics used in each model are indicated in the text.

For the classification of patches from the natural texture images we first separated the texture images into a training and a testing set, randomly assigning 319 texture images to each. Then for each texture image we generated all the unique combinations of pairs of patches for the matched condition (6 combinations). Then we generated pairs of patches from different textures (randomly sorted) within each image set, generating 10 pairs of these for each texture. This procedure generated 1914 matched pairs of patches and 1595 unmatched pairs of patches for each the training set and the testing set.

For the classification of patches from natural scenes, we randomly sorted the images into a training set of 332 images and a testing set of 84. We then generated the pairs of patches as described above, producing 2688 pairs of patches (1408 matched paris and 1280 unmatched). On average, there were 2150 pairs of patches in the training set, and 537 pairs in the testing set (there is some variability due to the image sorting, since not all images had the same number of segments).

We repeated the random sorting of training and testing set 20 times for each model. In the figures, we show the results for the model trained with each sorting, as well as the average performance.

### 2.7) Identifying pairs with useful HOS

To identify the pairs of patches where HOS improved segmentation (referred to as pairs with useful HOS for brevity), we first split the number of images in the dataset (either for BSD or for the textures) into 10 non-overlapping subsets, to be used as testing sets separately. Then, we iterated through all the 10 subsets of patches, training a segmentation model on the image pairs that did not belong to the testing subset, and then testing the model on the subset. For each subset we trained both a model using spectral statistics alone, and a model using HOS alone. Then, for each pair in the testing set, we compared the outputs of the two models, and we labeled all pairs of patches that were incorrectly classified by spectral statistics but correctly classified by HOS as having useful HOS. We repeated this procedure for all testing sets, obtaining a label for each pair of patches. The same procedure was performed comparing the model with spectral statistics alone to the model with both spectral and HOS.

Note that the size of the train and test sets used here are different from the main segmentation task. As described above, for the main segmentation task, when using the texture dataset half of the textures went into the training set, and when using natural images, 20% of the images went into the training set. Here, in both cases 90% of the dataset went into training for each model. This means that for the texture dataset, 6314 pairs of textures were used for training, and 704 for testing in each iteration. For the natural scenes dataset, on average 2419 pairs went into the training set and 269 into the testing set for each iteration.

Then, we again split the dataset into 10 subsets, and for each subset, we trained a linear classifier on the rest of the patches to identify whether the pairs had useful HOS or not, using as inputs for this task the spectral and HOS. This way, for each pair of patches we obtained a ground-truth label indicating whether it had useful HOS, as observed in the segmentation models, and the output of a classifier labeling it as having useful HOS or not.

### 2.8) Data and code availability

All the analysis code used in this work is available at https://git.io/JJNyr.

## 3) Results

We used a texture discrimination task to quantify the contribution of different image statistics to texture-based segmentation (**Figure 1**). Specifically, the texture statistics of two image patches are given as input, and a classifier indicates whether these two patches belong to the same image segment or not (matched and unmatched pairs of patches respectively). We considered different groups of image statistics of the PS texture model: marginal pixel statistics, Fourier power spectrum (spectral statistics), and higher-order statistics (HOS) (see Methods for further detail). To quantify the contribution of these statistics, we trained different models using the absolute difference between the values of these statistics and compared their performance at the task.

### 3.1) Differences in spectral statistics and HOS are redundant in natural images

We first studied the correlation between differences in spectral statistics and HOS across segments, as a basic estimate of redundancy **Figure 2** shows, for 2688 pairs of neighboring image patches sampled from 416 natural scenes (BSD, (Martin et al., 2001), the angle between their vectors of spectral statistics (which measures how different the spectral statistics are between the two patches), and the angle between their HOS statistics. We found a strong positive correlation between the spectral statistics angles and the HOS angles (**Figure 2**; Pearson correlation = 0.55, p = 2e-16, CI = [0.52-0.58]) for neighboring patches, suggesting a high redundancy between these statistics.

Besides the correlation, which indicates overall redundancy, a more relevant question is how much information the differences in the individual spectral statistics and HOS provide for the task of segmentation. To quantify this we next measured how the use of the spectral statistics difference compares to both the use of HOS, and to the combination of spectral statistics and HOS for segmentation.

### 3.2) Spectral statistics and HOS are redundant for natural scene segmentation

To test the information in the different sets of statistics and in their combination for image segmentation, we trained a linear classifier on the individual statistics of the PS texture model using ridge regression to solve a segmentation task (see Methods). We used the same 2688 pairs of natural scene patches as in **Figure 2**. We performed 20 repetitions of the task by randomly separating the image set into 332 images for the training set (an average of 2150 pairs of segments) and 84 for the testing set (an average of 537 segments) for each repetition.

**Figure 3B** shows that adding the spectral statistics to the marginal pixel statistics improved performance, albeit modestly. The combination of pixel and HOS performed better than pixel and spectral statistics, although the difference was small, and spectral statistics performed slightly better than HOS when both were used without pixel statistics. Finally, combining spectral and HOS led to an improvement in segmentation performance (both with and without pixel statistics), although the improvement was modest.

**Figure 3.**
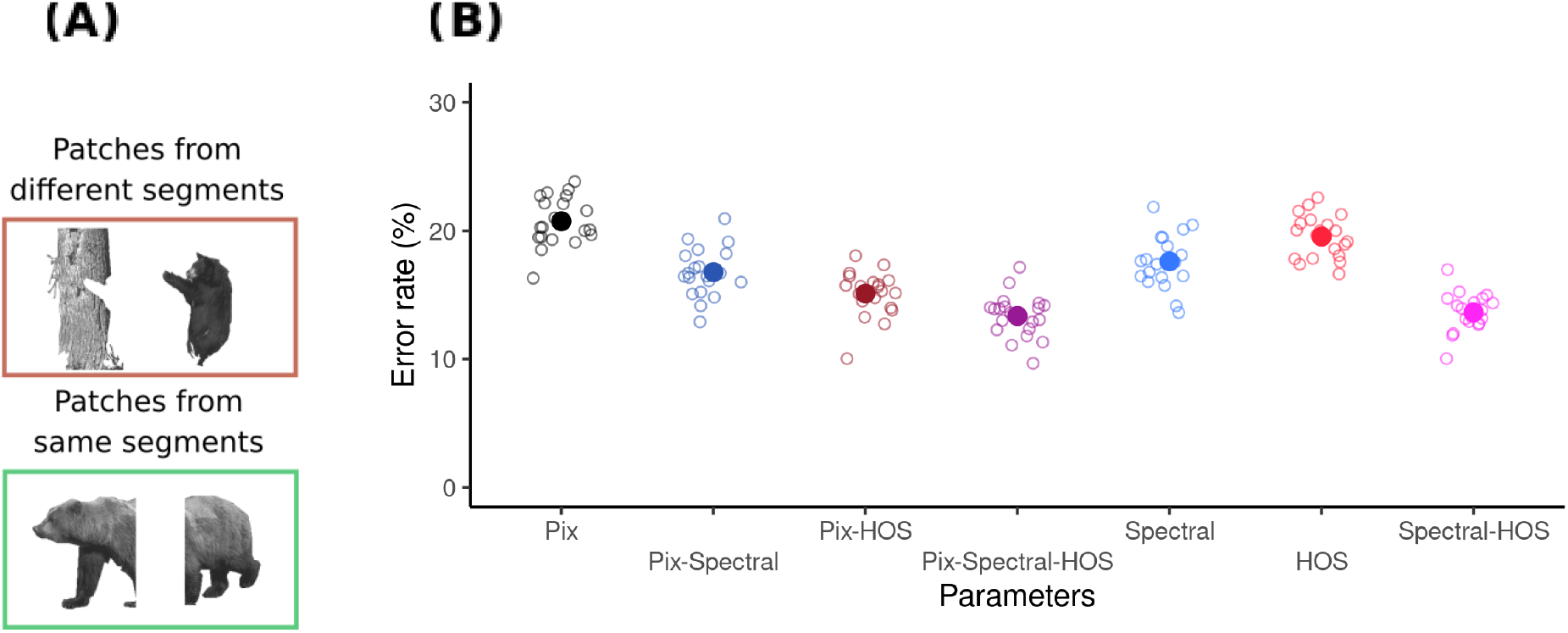
(A) Example of pairs of image patches used in the task. Top: Patches belong to the different segments. Bottom: Patches belong to the same segment. (B) Performance of a linear model in classifying pairs of image patches as belonging to the same or to different image segments. Models using different subsets of statistics from the PS model are shown. The empty circles show model performance for the 20 individual models trained and evaluated with different splits of training and testing data sets. The filled circles show the mean performance across splits.

These results show that the segmentation performance of the model using spectral statistics is high, and that it improves only modestly when adding HOS, even though HOS alone also achieved high performance. This supports the idea that the HOS of the PS model are highly redundant with the spectral statistics for segmenting natural images. We observed similar results with a non-linear decoder (i.e. a neural network), showing that our findings do not simply reflect a limitation of the linear decoder (**Tables S1, S2**, **Figure S1**).

We reasoned that our results could be influenced by differences in dimensionality: the spectral statistics of the Portilla-Simoncelli model are 4 times less numerous than the HOS (137 statistics and 552 statistics respectively, with our selected number of orientations, scales and neighborhood size), and we found similar ratios for their intrinsic dimensionality (**Table S3, S4**). To test this possibility, we used PCA to match the dimensionality of the spectral and HOS statistics, and we found similar results to **Figure 3B**(**Table S5**). Thus, despite the HOS being more numerous, when using subspaces with the same dimensionality, spectral and HOS statistics still perform similarly on segmentation.

### 3.3) Spectral statistics and HOS are redundant for natural texture segmentation

Next, we asked whether the contributions of these sets of statistics to the more specific task of segmenting natural textures, is similar to what we found for natural scenes. This is important because in the BSD, natural scenes have been segmented manually by human observers who likely used several other cues, in addition to texture, to determine segmentation. The use of these other cues for segmenting and grouping image patches in scenes may influence the observed distribution of texture features across and within segments. Thus, to better understand the contribution of HOS to natural texture segmentation, we next repeated the analysis for a texture segmentation task, using pairs of patches that were obtained from natural texture images (Brodatz, 1966; Lazebnik et al., 2005; *Salzburg Texture Image Database (STex)*, n.d.) (**Figure 4A**).

**Figure 4.**
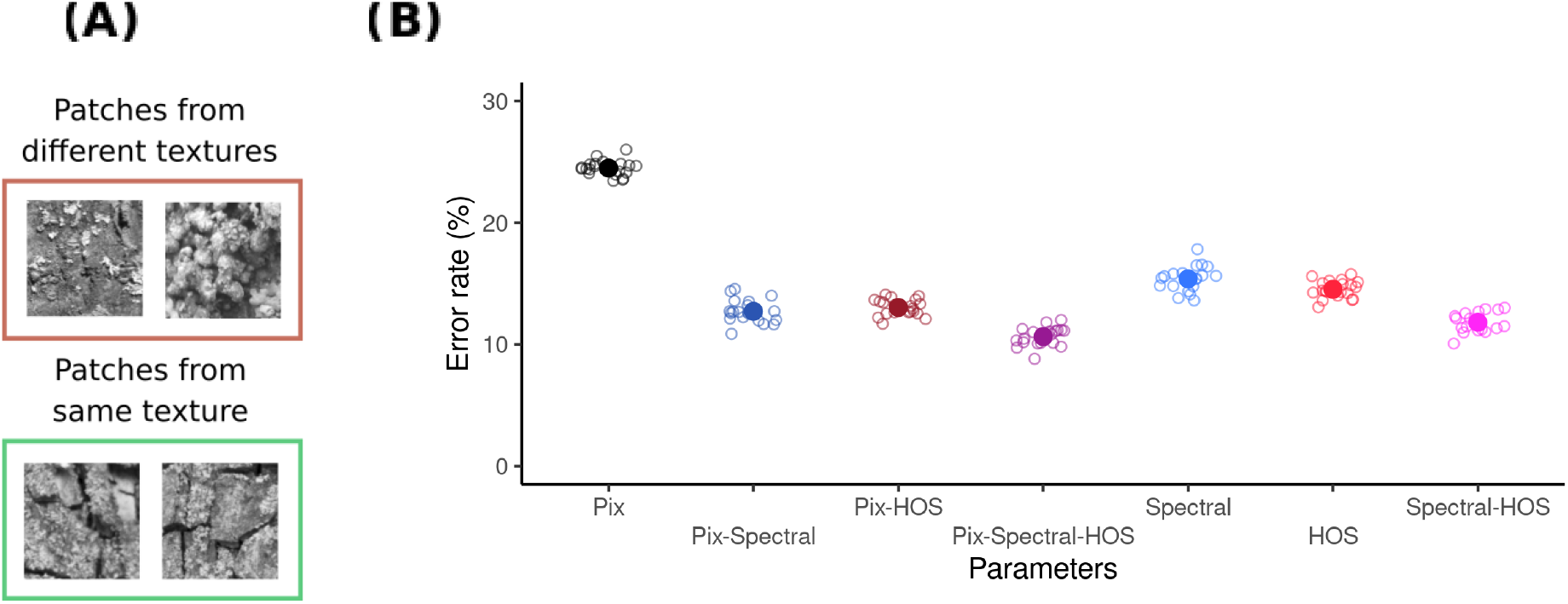
(A) Example of pairs of texture patches used in the task. Top: Patches belong to the different textures. Bottom: Patches belong to the same texture. (B) Performance of a linear model classifying pairs of texture patches as belonging to the same or to different textures. Models using different subsets of the PS statistics are shown. Same conventions as in Fig. 3B.

We trained the model on 3509 pairs of texture patches using ridge regression (see Methods), and then tested it on 3509 pairs of patches sampled from a different set of natural textures. We repeated this procedure 20 times, resampling the training and test sets.

We observed that segmentation performance for natural textures was in general higher than for the natural scenes (**Figure 4B**). Also, segmentation improved substantially when we added the spectral statistics to the pixel statistics. As with natural images, we observed that using pixel statistics and HOS led to similar performance than using pixel and spectral statistics. Using both the spectral statistics and HOS also improved performance over using the spectral statistics alone, although modestly.

These results indicate that spectral and HOS are redundant for the task of natural texture-segmentation. We note that although the results for texture segmentation are similar to those for scene segmentation, models trained on one dataset generalize poorly to the other (**Table S6**), supporting the hypothesis that different texture features may be required for the two tasks.

### 3.4) Images with useful HOS are difficult to identify

In our experimental work (Herrera-Esposito et al., 2021), we observed considerable between-texture variability in the effect of HOS on segmentation. Similarly, previous work (Freeman et al., 2013; Okazawa et al., 2015, 2017; Ziemba et al., 2016) showed that different synthetic PS textures lead to different perceptual and neural discriminability. Therefore, we considered the possibility that, although combining spectral and HOS leads to a modest performance improvement overall, HOS could be particularly useful for some subset of images.

To analyze this possibility, we identified pairs of natural scene patches where classification was better when using HOS. Specifically, we compared for each pair of patches the classification with spectral statistics alone to HOS alone (we obtained similar results when using spectral and HOS together, data not shown).

Overall, 10% of the full set of pairs were better classified by HOS than by spectral statistics, with similar proportions for pairs belonging to the same and to different segments (data not shown). Conversely, 12.5% of the pairs were better classified by spectral statistics than by HOS. In particular, 59% of the pairs misclassified by spectral statistics were correctly classified by HOS, confirming that HOS are useful for some images. In addition, if segmentation by spectral and HOS were independent, the error rates for HOS should be the same in the complete dataset (19.3%) as in the subset misclassified by spectral statistics. Instead, 41% of the pairs misclassified by spectral statistics were also misclassified by HOS, reflecting the redundancy in the responses of the two sets of statistics.

We next tested whether this subset of images with more useful HOS can be identified from their statistics, which would be required for the visual system to use HOS more strongly in these cases. For this, we relabeled the pairs of patches to indicate whether segmentation was better when using HOS or not, and we then trained a new linear classifier on these labels, using spectral and HOS together as predictors (see Methods).

The confusion matrix (**Table 1**) shows that the classifier performed better than chance, (p < 2e-16, McNemar’s test), which indicates that there is some consistent difference between pairs where HOS improve segmentation and those where it does not. However, due to the imbalance between the classes, we observe a low overall accuracy of 68%, that is lower than the accuracy obtained by classifying all pairs as not being improved by HOS. In line with these results, visual inspection of the pairs of patches better classified by HOS does not show obvious patterns that distinguish them from other pairs (**Figure 5A**).

**Table 1.** Classification of pairs of natural image patches as being better segmented by HOS or not. The columns of the table indicate the observed ground truth label, of whether a pair of patches was better labeled by HOS than by spectral statistics or not. The rows indicate the label for the pairs of patches predicted by a linear classifier. Each cell in the table shows the number of pairs for each combination of true and predicted labels.

|  | Label: No HOS improvement | Label: HOS improvement |  |
| --- | --- | --- | --- |
| Predicted: No HOS improvement | 1707 | 160 | 1867 |
| Predicted: HOS improvement | 711 | 110 | 821 |
|  | 2418 | 270 |  |

**Figure 5.**
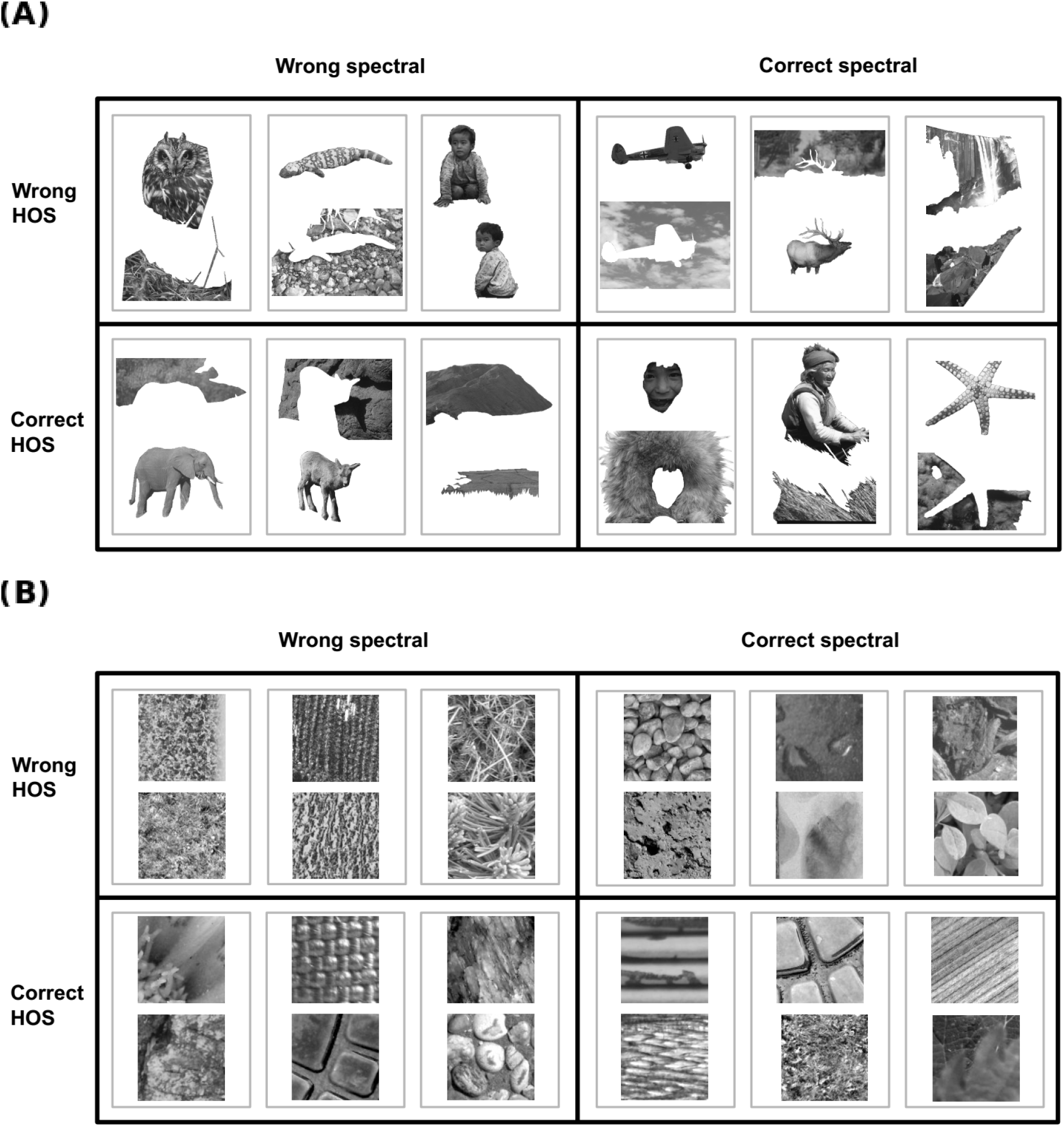
**(A)** Examples of pairs of natural scene patches with different classification outcomes for both spectral and HOS. Only pairs extracted from different segments are shown. Smaller gray boxes group the patches that form a pair. Larger black boxes indicate the classification outcome for both spectral and HOS. **(B)** Same as **(A)** but for natural textures.

We obtained similar results for texture segmentation. We found that 8.7% of the texture pairs were better classified by HOS than by spectral statistics, and that 8.4% were better classified by spectral statistics than by HOS. Of the pairs misclassified by spectral statistics, 58% of which were also misclassified by HOS, showing again redundancy in their responses. A classifier trained to identify the patches with HOS improvement, as described for BSD above, had a performance better than chance (p < 2e-16, McNemar’s test, **Table 2**), but with a low accuracy of 67%. **Figure 5B** shows some example pairs of textures with different classification outcomes for spectral and HOS (more example pairs can be found together in the open repository with the analysis code).

**Table 2.** Classification of pairs of texture image patches as being better segmented by HOS or not. Same conventions as **Table 1**

|  | Label: No HOS improvement | Label: HOS improvement |  |
| --- | --- | --- | --- |
| Predicted: No HOS improvement | 4324 | 271 | 4595 |
| Predicted: HOS improvement | 2087 | 336 | 2423 |
|  | 6411 | 607 |  |

In sum, the misclassifications of spectral and HOS showed redundancy in both natural scenes and textures, but a subset of the pairs of patches was better classified by HOS. Nonetheless, the pairs better classified by HOS were not accurately identified by a linear classifier (for further analysis on the causes of the low accuracy see Supplementary section **S4**). We also obtained similar results when using a procedure to reduce possible labeling noise, in which we trained to separate models on non-overlapping subsets of the training set, and required that HOS be better than spectral statistics in both training sets for a given pair, in order to label that pair as having useful HOS (results not shown).

Furthermore, we compared the predictions from these models to our previous experimental results (Herrera-Esposito et al., 2021), to test whether the observed experimental variability between pairs of textures correlates with the estimated usefulness of the HOS. We did not observe any clear agreement between the two that could be suggestive of fine-tuning to use HOS in informative cases (**Figure S2**, although the low number of textures and several other caveats demand caution when interpreting these results, see the Supplementary section **S5**).

### 3.5) Subsets of HOS contribute differently to segmentation

Besides the results from previous studies mentioned above, showing that different PS textures drive mid-level visual areas and perception to different degrees, the same line of research has also identified specific subsets of HOS as driving perception (Freeman et al., 2013; Hermundstad et al., 2014; Tesileanu et al., 2020; Victor et al., 2013) and physiology (Freeman et al., 2013; Okazawa et al., 2015, 2017; Yu et al., 2015) to different degrees. Also, this has been shown to follow natural image statistics (Hermundstad et al., 2014; Tesileanu et al., 2020; Yu et al., 2015). Therefore, we next wondered whether different subsets of HOS would show varying degrees of usefulness in our segmentation task.

To determine the relevance of different subsets of HOS to our segmentation model, we divided the HOS into the four subsets used in the PS model (Portilla & Simoncelli, 2000): energy correlations across space, energy correlations across scale, energy correlations across orientation, and phase correlations across scale (also called linear correlations across scale in some studies). We then tested the performance of different combinations of these subsets of HOS, with and without spectral statistics, in the segmentation task.

**Figure 6** shows the performance of the segmentation models (top row, natural scenes; bottom row, natural textures) using different subsets of HOS. All the subsets of HOS alone had considerably worse performance than spectral statistics (indicated by the dashed blue line, **Figure 6A**) for natural scene segmentation. Adding each subset of HOS to spectral statistics did not reach the performance of the full model (purple line, **Figure 6B**). Similarly removing each subset of HOS never decreased performance to the level of spectral statistics alone (**Figure 6C**). In most cases, spatial correlations were the most useful subset of HOS. Also, orientation and phase statistics were the least useful when considered alone and together with spectral statistics, but phase statistics gained in relevance when removing the subsets of HOS from the full model.

**Figure 6.**
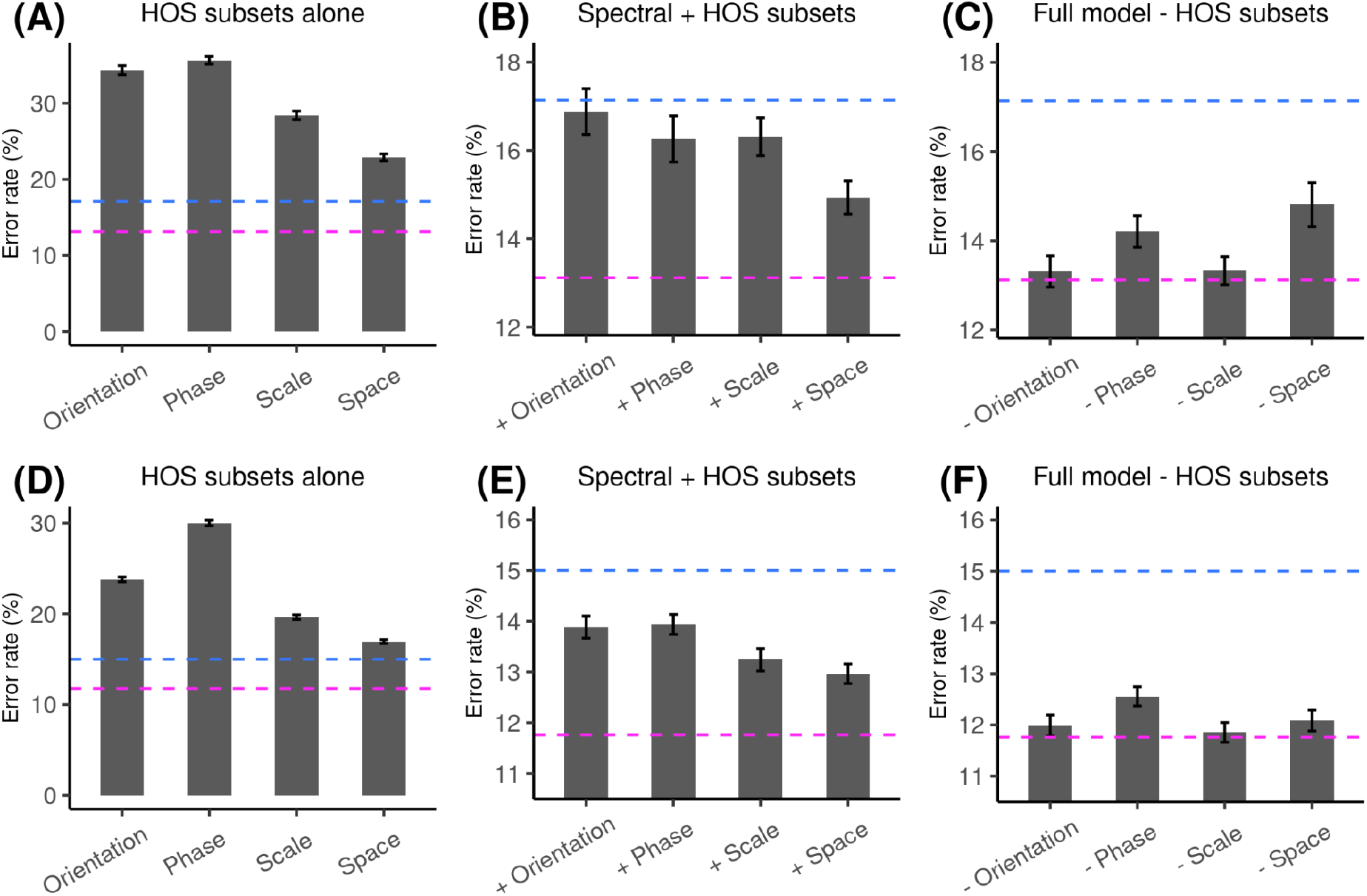
Performance at the segmentation task using different subsets of HOS. Each grey bar shows the average over the outcome of 50 different models trained on random splittings of the data into train and test set. The error bars show the 95% confidence interval of the mean. The dashed horizontal blue line shows the performance for the model using only spectral statistics, and the red line shows the performance of the model with spectral and all HOS. **(A)** Segmentation performance for the BSD using subsets of HOS alone. **(B)** Segmentation performance for the BSD using subsets of HOS together with spectral statistics. **(C)** Segmentation performance for the BSD using the model containing spectral statistics and all subsets of HOS except those indicated in the horizontal axis. **(D)**, **(E)** and **(F)**, same as **(A)**, **(B)** and **(C)** but for the texture segmentation dataset.

Results for textures (**Figure 6D-F**) were similar to those for natural scenes, except that adding orientation correlations improved performance more markedly, and removing spatial correlations did not reduce performance.

These analyses confirm that different HOS subsets have different usefulness for segmenting natural scenes and natural textures. Spatial correlations seem to be, in general, the most informative subset of HOS, and in most cases they were followed by correlations across scale. Phase correlations allowed for improved performance when combined with spectral statistics, and they had a considerable effect when removed from the full model, indicating that they contain useful information that is not redundant with the rest of the HOS. Correlations across orientation were generally among the least useful for segmentation when considering models containing spectral statistics.

## 4) Discussion

We have studied the importance of different image statistics, namely the spectral statistics and HOS of the PS texture model, for segmenting natural textures and images. First, we showed that there is a strong correlation between the difference in spectral statistics and the difference in HOS for pairs of neighboring patches in natural scenes (**Figure 2**). Then, using segmentation tasks with either natural scenes segmented by human observers or natural textures, we showed that using either the spectral statistics alone or the HOS alone were enough to solve the task with high accuracy, indicating they are both reliable cues for segmentation. Importantly, combining both together produced modest improvements, for both linear and non-linear classifiers (**Figures 2, 3, S1, Table S2**). These results indicate a strong redundancy between spectral statistics and HOS specifically in the context of image segmentation, and seem to rule out the alternatives that HOS cues for segmentation are either unreliable or largely independent from spectral statistics.

In a recent study on human texture perception, we reported that differences in the HOS of the PS model between adjacent textures in peripheral vision produced only weak segmentation when the textures had matched spectral statistics (Herrera-Esposito et al., 2021). In another related study (Balas, 2008), the author observed that human texture similarity judgements were better predicted by the power spectrum of the textures alone, than by the entire set of PS statistics. These results are of particular interest because these statistics have high perceptual relevance (Balas, 2006; Freeman et al., 2013; Freeman & Simoncelli, 2011; Portilla & Simoncelli, 2000; Wallis et al., 2017) and drive neural activity in mid-level visual areas (Freeman et al., 2013; Okazawa et al., 2015, 2017; Ziemba et al., 2016). Furthermore, these statistics are related to the second processing stage in the SS model of peripheral vision (Balas et al., 2009; Freeman et al., 2013; Freeman & Simoncelli, 2011; Rosenholtz, 2016), of which segmentation has been argued to be an important missing component (Doerig et al., 2019; Herrera-Esposito et al., 2021; Herzog et al., 2015; Manassi et al., 2012, 2013; Wallis et al., 2019), making their role in segmentation an essential aspect for the further development of this model. In the present work we expand on our previous results showing that the small effect observed for these HOS on perceptual segmentation may be related to their redundancy with spectral statistics in natural images for the task of image segmentation (**Figure 1**), since they may not add much to the initial segmentation process based on the power-spectrum representation in V1 (V. A. Lamme, 1995; Landy & Bergen, 1991; Z. Li, 2002; Nothdurft et al., 2000; Victor et al., 2017).

In line with this argument, previous work showed that the higher variability in second-order pixel statistics in natural images as compared to third and fourth-order pixel statistics matched their perceptual saliency (Hermundstad et al., 2014; Tesileanu et al., 2020; Tkacik et al., 2010). Nonetheless, besides using a different kind of texture than the ones presented in this work and our experimental study (Herrera-Esposito et al., 2021), the computational analysis of image statistics in these previous studies was performed in the context of efficient coding, rather than the specific perceptual task of image segmentation. Different tasks may rely on different texture properties (Victor et al., 2017), which can explain why the HOS of the PS model are simultaneously very important for texture perception (Balas, 2006; Portilla & Simoncelli, 2000; Wallis et al., 2017) but maybe less so for segmentation. A variation of this idea is also espoused in those previous studies on natural texture statistics (Hermundstad et al., 2014; Yu et al., 2015), where it is noted that the sensory periphery (i.e. the retina) and the cortex face different constraints and goals that lead to different coding regimes. In this sense, the present work is a contribution to the growing efforts of comparing perceptual systems to model observers performing sophisticated tasks on natural images (Burge, 2020).

But although there is a high redundancy between the spectral statistics and the HOS for image segmentation, the observation that HOS are a reliable segmentation cue and that they can improve texture segmentation, raises the question of why the spectral statistics and not the HOS are used as a strong segmentation cue, as shown in peripheral vision (Hermundstad et al., 2014; Herrera-Esposito et al., 2021; Victor et al., 2013). One possible explanation to this regard is the constraint in resources that makes information processing by the visual system a balance of costs and benefits. While the HOS improved model performance, they did so only modestly and one could hypothesize that the cost of using these HOS would lead the visual system to use the spectral statistics as the main segmentation cue, with these HOS being a secondary or null segmentation cue.

Nonetheless, a softer version of the hypothesis is that the HOS of the PS model are particularly useful in some scenarios (i.e. specific kinds of images), and that the visual system is fine-tuned to rely on HOS in these cases. As mentioned previously, this could be in line with our previous experimental work on perceptual human segmentation (Herrera-Esposito et al., 2021), as well as with previous physiological and perceptual work studying PS textures (Freeman et al., 2013; Okazawa et al., 2015, 2017; Ziemba et al., 2016). Since this fine-tuning would rely on the ability to identify which images have useful HOS for segmentation, here we tested whether a linear model could identify the pairs of patches where HOS improved segmentation over spectral statistics. We found that these pairs of patches were difficult to identify (**Tables 1, 2**), suggesting that this kind of fine-tuning may be difficult to achieve in practice. Furthermore, when comparing the predictions from the models to our experimental results reported in (Herrera-Esposito et al., 2021) we did not find any agreement between models and experiment that could suggest such a fine tuning (**Figure S2**, although this analysis is preliminary due to the little experimental data available, and should be interpreted with caution).

Although the pairs of patches with useful HOS could not be clearly identified, we did find that some subsets of HOS are more useful than others for segmentation, both in isolation and in combination with spectral statistics (**Figure 6**). Mainly, we observed that when considering the subsets of HOS separately, spatial and scale energy correlations were generally the ones with best segmentation performance (**Figures 6A, 6B, 6D, 6E**). This finding agrees with previous studies showing that scale and spatial energy correlations explain the most variance in the variability of perceptual sensitivity between PS textures (Freeman et al., 2013), and in V4 neurons ability to discriminate PS textures from noise (Okazawa et al., 2015).

The agreement between our analysis and these previous results may reflect a fine tuning of the visual system to the usefulness of the different subsets of HOS, which is captured in our analysis of segmentation. But this agreement does not necessarily mean that the visual system is tuned to use these HOS specifically for segmentation. One alternative is that these HOS are the most informative ones in general, and the visual system is tuned to use them for other tasks as well. In relation to this, (Okazawa et al., 2015) reported that energy correlations across space are the HOS with highest performance in a texture classification task, which they propose as a possible explanation to their physiology results. Furthermore, the ordering of HOS relevance may also depend on what ranking criterion is used, requiring careful comparisons across tasks and methods. For example, we observed a different ordering of the relevance of HOS subsets when performing segmentation alone than when in the context of the full model (**Figures 6C, 6F**). In line with this, the ranking of HOS subsets relevance obtained from analyzing their contribution to discriminability of textures from spectrally matched noise in V4 is somewhat different from the ranking obtained for explaining V4 responses to textures in general (Okazawa et al., 2015, 2017). Therefore, more work is needed to understand how the information different HOS subsets carry for segmentation in natural images, relates to their use by the visual system.

In conclusion, the results presented here, show that spectral statistics and the HOS of the PS model have a strong redundancy for natural scene and texture segmentation, which coupled with resource constraints may explain the weak effect of these HOS in human segmentation (Herrera-Esposito et al., 2021). This also suggests that segmentation based on the HOS of the PS model may not be crucial to future extensions of the SS model of peripheral vision, but rather that existing models of segmentation based on the outputs of V1-like oriented filters that respond to spectral statistics (Bergen & Landy, 1991; Bhatt et al., 2007; Z. Li, 2002) may be enough to considerably expand its explanatory power. Nonetheless, there are some important caveats that need to be considered.

One important caveat is that the redundancy between spectral and HOS reported here is compatible with either of them taking a secondary role. Although there is plenty of evidence showing the primacy of spectral statistics over HOS in texture segmentation, these studies have been mostly done in the peripheral visual field. Therefore, it is not clear whether the same holds for central vision, and our results here do not necessarily mean that HOS take a secondary role there. Furthermore, our main line of argument rests on the potential cost of using HOS for segmentation, and the role of resource constraints in the brain. Given that resources are much more constrained in the periphery than in central vision, our line of thought is compatible with a stronger role of HOS for segmentation in central vision.

Another important caveat is that we only considered a specific set of HOS, the ones in the PS model. While the HOS of the PS model capture to a considerable extent the perceptual quality of natural textures, they sometimes fail to fully reproduce their structure (Portilla & Simoncelli, 2000). Therefore, other HOS not present in the PS model are important for texture perception, and it is possible that they contribute more strongly to segmentation, both in humans and in segmentation models. One example is the correlations between the features of mid-level layers in deep neural networks, which have been shown to capture the visual appearance of many textures (Gatys et al., 2015), and that allow for good performance in image segmentation (Vacher & Coen-Cagli, 2019).

On the other hand, we also did not consider other low-level segmentation (or saliency) cues that are represented in V1, such as color, binocular disparity and motion (Braddick, 1993; A. Li & Lennie, 2001; Møller & Hurlbert, 1996; Nakayama et al., 1989; Saarela & Landy, 2012). A more general version of our main hypothesis could be that, for segmentation, the HOS of the PS model are redundant with the cues available in V1 in general, and not only with spectral statistics. This alternative hypothesis would be more in line with the proposal that there is a bottom-up saliency or segmentation map in V1 based on these features (Z. Li, 2002; Zhang et al., 2012; Zhaoping, 2019). Therefore, it is possible that by ignoring these other early segmentation cues, we overestimated the contribution of HOS to bottom-up segmentation. This more general hypothesis may also explain why, being redundancy a mutual relationship where either kind of statistics could be used, it is the HOS that adopt a secondary role.

The last important consideration is that we used a region-based texture segmentation task (i.e. using the properties of two image regions to decide whether they belong to the same segment), but segmentation may also proceed through processes based on identifying texture-defined edges (Giora & Casco, 2007; Landy, 2013; Machilsen & Wagemans, 2011; Rosenholtz, 2014). This other type of model may change some of the analysis regarding the possible roles of HOS. For example, the most popular edge-based texture segmentation model is the Filter-Rectify-Filter (FRF) model, which consists in a V1-like filtering with rectification, followed by a second filtering stage capable of detecting texture-defined edges (Landy, 2013). Depending on the non-linearity used in such models (among other possible modifications), they may be able to find edges defined by HOS discontinuities, and these models have been shown to correlate with human HOS-based segmentation in central vision in one study (Zavitz & Baker, 2014). It is interesting to note that such a segmentation process could show sensitivity to HOS, even though still operating directly on rectified V1 outputs, instead of operating on units that encode HOS directly, such as V2 neuron outputs. This means that a simple segmentation model operating over V1 outputs could still explain some effect of HOS such as those observed in our experimental work (Herrera-Esposito et al., 2021). Another important edge-based segmentation mechanism is the emergence of selectivity to texture borders by tuned contextual modulation, which can emphasize the response of neurons near texture edges. This mechanism is proposed to be an important mechanism for computing segmentation and saliency in this area (Z. Li, 1999, 2002; Nothdurft et al., 2000). It is difficult to anticipate how HOS may affect these complex mechanisms when they operate on V1-like outputs, but they could lead to effects on segmentation that are not captured by our region-based segmentation task. Also, in another previous study (Ziemba et al., 2018), it is shown that contextual modulation for textures in V2 neurons is tuned to the HOS of the PS model, which could also give rise to this kind of segmentation based on contextual-modulation within V2. Nonetheless, see also (Schmid & Victor, 2014) where V2 neurons responses to texture-defined edges are argued to be compatible with a filter-rectify-filter mechanism, and less so with this kind of contextual modulation mechanism.

## Supporting information

Supplementary_results

## Acknowledgments

We thank Ruth Rosenholtz for useful comments on previous work that motivated this analysis. Ruben Coen-Cagli was supported in part by National Institutes of Health grant EY031166. Daniel Herrera-Esposito was supported by the studentships from the Comisión Académica de Posgrados, UdelaR, Uruguay, and by travelships awarded by PEDECIBA, UdelaR, Uruguay, and CSIC, Uruguay.

