## Supplementary_results for "Redundancy between spectral and higher-order texture statistics for natural image segmentation"

### Index:

S1. Non-linear and linear decoding show similar redundancy……………………….p1
S2. Spectral and HOS still perform similarly when matched in dimensionality……. p4
S3. Models generalize poorly between scenes and texture datasets………………..p5
S4. Identification of pairs with useful HOS fails due to within-class heterogeneity…p6
S5. Low agreement between models of HOS fine-tuning and psychophysics……...p9

### S1) Non-linear and linear decoding show similar redundancy:

To test whether a non-linear decoder could extract further information from the statistics for segmentation, we analyzed whether the same patterns in performance were observed when neural networks were used to solve the natural image segmentation task.

We trained the neural networks using the Keras package for R (Francois Chollet, 2015) to classify the pairs of patches from the BSD, using the same training and testing division scheme as described for the linear classifier in the main text. We also normalized the differences in statistics of the pairs of the training set to have 0 mean and unit variance, and performed PCA on these values, keeping the principal components that explained 95% of the variance. This resulted in the number of principal components for each model shown in **Table S1**. Although we applied PCA simultaneously on all the statistics used by a given model, performance was similar when applying PCA on the subsets of statistics individually.

**(SP1)** To select the architecture and training regime of the network, we performed a search of parameters that maximized performance at the task. For this, we focused specifically on the performance of the model using pixel and HOS, which in preliminary analyses showed worse performance than the linear classifiers (suggesting overfitting of the nonlinear classifier). All networks had units with ReLu activation functions, and an output layer of 1 unit with a sigmoid activation function. Furthermore, all were trained using a binary cross entropy loss with the Adam gradient descent algorithm, with a batch size of 32. We tested 8 fully connected networks with the following architectures, with the hyphen separated numbers indicating the number of units in successive hidden layers in the network: 10, 30, 50, 10-2, 30-10, 50-20, 30-10-2, 50-20-5. Thus, we tested networks with 1 to 3 hidden layers, and different numbers of units. We also tested three different L1 regularization penalties: 0.001, 0.003 and 0.01. Furthermore, we trained the networks for a duration of 350 epochs. For each combination of parameters, 10 models were trained and tested on different random splits of the data into training and testing sets.

| **Statistics** | **Principal components (95% variance)** |
|---|---|
| Pixel | 8 |
| Pixe + Spectral | 45 |
| Pixel + HOS | 176 |
| Pixel + Spectral + HOS | 206 |
| Spectral | 38 |
| HOS | 169 |
| Spectral + HOS | 200 |

**(SP2) Table S1**. Number of principal components that retain 95% of the variance for each combination of statistics. The PCA was performed on all statistics in the indicated model together.

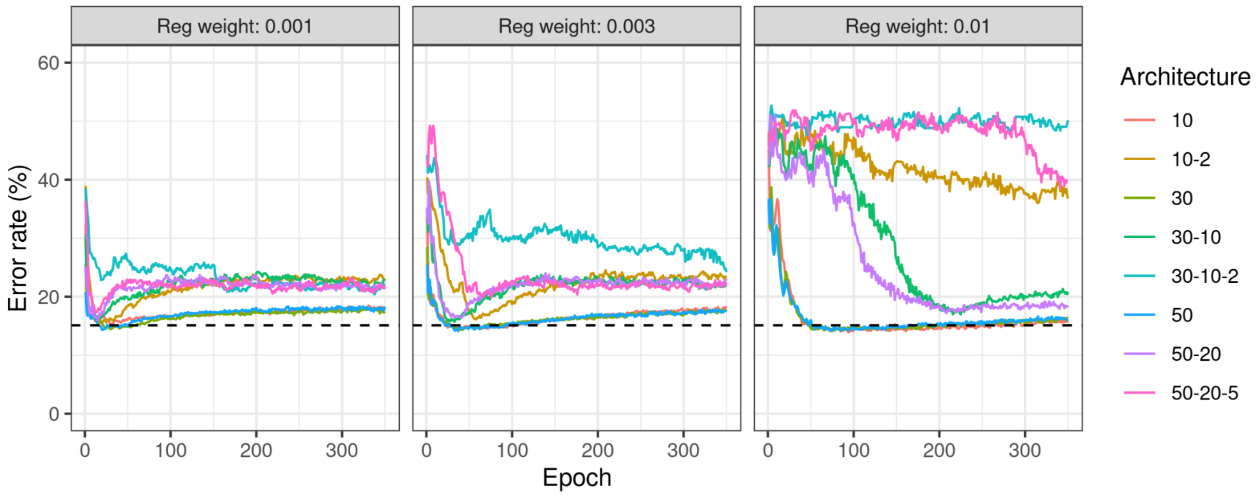


**Figure S1.** Segmentation performance during training for neural networks using HOS. Each line shows the average performance for 10 model instantiations using different train-validation set splits. The architecture of the model corresponding to each colored line is indicated in the legend to the right.

The horizontal dashed line shows the performance for the same set of statistics when using the linear model.

In **Figure S1** we see the performance throughout training for the different models using pixel and HOS. We observe that the networks that achieve the best performance are the simplest ones, consisting of only one hidden layer. Furthermore, these networks achieve best performance early in training, and then their performance deteriorates, probably due to overfitting. Also, we see that peak performance is similar to the performance of the linear model (indicated by the dashed horizontal line).

Therefore, following the results from the parameter exploration in **Figure S1**, we next tested a network with a single hidden layer of 30 units on all sets of statistics, using 80 epochs for training. The performance results shown in **Table S2** show that performance changes only slightly for all sets of statistics compared to the linear model.

| **Parameters** | **Linear model: % error rate (SD)** | **Neural network: % error rate (SD)** |
|---|---|---|
| Pixel | 20.7 (1.9) | 20.0 (1.9) |
| Pixel + Spectral | 16.8 (1.9) | 16.3 (1.9) |
| Pixel + HOS | 15.1 (1.8) | 14.4 (1.7) |
| Pixel + Spectral + HOS | 13.4 (1.7) | 13.0 (1.7) |
| Spectral | 17.6 (2.1) | 17.0 (1.5) |
| HOS | 19.5 (1.7) | 18.4 (1.7) |
| Spectral + HOS | 13.6 (1.5) | 13.0 (1.3) |

**Table S2.** Mean performance error in the segmentation task on natural image patches using each combination of texture statistics, with either the linear classifier shown in **Figure 3** in the main text, or using a neural network decoder with one hidden layer of 30 units. For each combination of statistics and each type of model, 20 instances of the model were trained with random splittings of the patches into train and test sets.

We note that although selecting the number of training epochs for which HOS performance was optimal may bias the resulting performances in favor of HOS, when analyzing the performance changes during training for spectral statistics and for the full model, the results were similar to those of HOS (data not shown).

### S2) Spectral and HOS still perform similarly when matched in dimensionality:

Although the number of HOS in the PS model (552) is considerably larger than the number of spectral statistics in the PS model (137), both sets of statistics have considerable redundancy. Thus, we analyzed whether this difference in the number of statistics is maintained when performing PCA on these subsets. In **Table S3** we show the number of principal components (PC) required to retain different amounts of variance in the Berkeley Segmentation Dataset (BSD). We observe that the number of components needed to retain a fixed amount of variance was around 4 to 5 times larger for HOS than for spectral statistics in all cases. In **Table S4** we also see that the performance of the linear segmentation model using the PC for different levels of retained variance maintains roughly the same patterns as when using the original statistics space.

| Statistics | Original number | 95% variance | 90% variance | 80% variance | 60% variance |
|---|---|---|---|---|---|
| Pixel | 16 | 8 | 6 | 5 | 2 |
| Spectral | 137 | 38 | 24 | 12 | 4 |
| HOS | 552 | 169 | 112 | 61 | 23 |

**Table S3.** Number of principal components needed to retain the indicated percentages of variance of the pairs of BSD, for each combination of statistics.

| Statistics | % error rate (all stats) | % error rate (95% variance PC) | % error rate (80% variance PC) | % error rate (60% variance PC) |
|---|---|---|---|---|
| Pixel | 20.7 | 20.8 | 23.0 | 29.7 |
| Pixel + Spectral | 16.8 | 17.5 | 18.3 | 19.9 |
| Pixel + HOS | 15.1 | 16.3 | 17.2 | 20.6 |
| Pixel + Spectral + HOS | 13.4 | 13.8 | 14.7 | 16.5 |
| Spectral | 17.6 | 19.1 | 21.5 | 22.2 |
| HOS | 19.5 | 20.2 | 21.1 | 24.1 |
| Spectral + HOS | 13.6 | 14.5 | 16.4 | 17.9 |

**Table S4.** Segmentation performance in natural images using the PCs of each subset of statistics that retain the indicated amount of variance. Error rates show the average of 20 models trained on different random train-test splits.

Although this could be initially interpreted as spectral statistics offering similar performance with fewer parameters, when setting the number of PCs of HOS to be the same as for spectral statistics, the trends in performance across models do not change much, as shown in **Table S5.** This indicates that, at least roughly, spectral and HOS can perform segmentation to similar performances using similar numbers of dimensions.

| **Statistics** | **% error rate (all stats)** | **% error rate (95% variance PC)** | **% error rate (80% variance PC)** | **% error rate (60% variance PC)** |
|---|---|---|---|---|
| Pixel | 20.7 | 20.8 | 23.0 | 29.7 |
| Pixel + Spectral | 16.8 | 17.5 | 18.3 | 19.9 |
| Pixel + HOS | 15.1 | 17.4 | 20.0 | 23.1 |
| Pixel + Spectral + HOS | 13.4 | 13.9 | 15.4 | 16.5 |
| Spectral | 17.6 | 19.1 | 21.5 | 22.2 |
| HOS | 19.5 | 22.2 | 25.0 | 27.3 |
| Spectral + HOS | 13.6 | 15.2 | 17.5 | 18.5 |

**Table S5.** Segmentation performance in BSD using the components of spectral and pixel statistics that retain the indicated amount of variance, but fixing HOS to have the same number of PC as spectral statistics. Error rates show the average of 20 models trained on different random train-test splits.

### S3) Models generalize poorly between scenes and texture datasets:

We hypothesized that due to the possible differences in the task of segmenting natural images (where cues other than texture are used to generate the labels) and texture segmentation, the results from scene segmentation may not be fully representative of the more specific process of texture segmentation in natural images. Since we found that our comparisons between the different kinds of statistics were qualitatively similar between the two, we wondered whether texture statistics may be useful in the same way for the two tasks. To answer this question, we tested whether the models trained in one dataset (i.e. BSD or our natural texture dataset) generalized to the other. For this, we used the same procedure for splitting each dataset into testing and training sets as described in the main text, but trained in the training set of one dataset, and tested in the other.

We observe in **Table S6** the performance results for these cross-trained models was considerably worse than for the models trained in the same dataset as they are tested (i.e. **Figure 3** and **Figure 4** in the main text), indicating that generalizability between the two datasets was poor.

| **Statistics** | **% error rate in natural images (trained with textures)** | **% error rate natural Images (trained with natural images)** | **% error rate in textures (trained in natural images)** | **% error rate in textures (trained in textures)** |
|---|---|---|---|---|
| Pixel | 35.6 | 20.7 | 37.9 | 24.5 |
| Pixel + Spectral | 28.8 | 16.8 | 21.4 | 12.7 |
| Pixel + HOS | 36.0 | 15.1 | 49.1 | 13.0 |
| Pixel + Spectral + HOS | 32.3 | 13.4 | 45.6 | 10.6 |
| Spectral | 27.6 | 17.6 | 20.2 | 15.4 |
| HOS | 33.4 | 19.5 | 50.6 | 14.5 |
| Spectral + HOS | 30.6 | 13.6 | 45.8 | 11.8 |

**Table S6. Segmentation performance of models trained in one of the datasets and tested in the other. Error rates show the average of 20 models trained on different random train-test splits.**

### S4) Identification of pairs with useful HOS fails due to within-class heterogeneity:

We wondered whether we could interpret some of the reasons behind the low accuracy of the model trained to detect useful HOS. For this, we noted that there are some important characteristics of the data that is being fitted that could be related to this behavior. These characteristics are:

1) The “No HOS improvement” class is heterogeneous, containing pairs where: a) spectral and HOS are wrong, b) spectral statistics are correct, HOS are wrong, and c) spectral and HOS are both correct.

2) Each class in the dataset (“No HOS improvement”, “HOS improvement”) also has heterogeneity because they contain pairs where the segmentation ground truth is “matched” and where it is “unmatched”. That is, the improvement in classification by using HOS can be either because HOS show a small difference between the patches that favors no segmentation (when the ground truth is “matched”) or because HOS show a large difference that favors segmenting the patches (when the pair is “unmatched”).

3) The dataset is highly imbalanced, with only 10% of the pairs of patches being "improved by HOS".

These properties of the dataset are due to the nature of the problem. 1) is because we want to find examples where HOS improves segmentation, 2) is because in order to have a classifier that tells us whether to use HOS or spectral statistics, and that is useful, we would want it to work without knowing the ground truth of the segmentation task, and 3) is due to how the classes are constructed, following 1). We hypothesize that it is these properties of the dataset (which follow from the nature of the problem) that make classification of HOS usefulness a difficult task.

Therefore, we propose that a control case in which our method for identifying pairs with useful HOS is expected to show high performance is when we apply the method to a subset of the data without characteristics 1), 2) and 3). To test this, we therefore trained the classifier on a subset of the data which has the following characteristics:

1) For the "Not improved" class we only use pairs of patches where spectral statistics are correct and HOS are incorrect. That is, we discard the pairs of patches where both statistics agree.

2) We only use pairs of patches where the ground truth segmentation is "unmatched". This way, HOS usefulness is because it favors segmentation.

Because of the criteria described above, this subset of the data is also much more balanced across classes, due to the removal of datapoints from the "Not improved" class.

When performing the same procedure as in the manuscript for this subset of data we obtain an accuracy of 86%, much higher than for the complete dataset (**Table S7**).

| | **Label: No HOS improvement** | **Label: HOS improvement** | |
|---|---|---|---|
| **Predicted: No HOS improvement** | 157 | 28 | 185 |
| **Predicted: HOS improvement** | 16 | 118 | 134 |
| | 173 | 146 | |

**Table S7.** Classification of pairs of texture image patches as being better segmented by HOS or not, where the pairs of patches with agreement between spectral statistics and HOS and pairs with "matched" ground truth are not included in training and testing.

Interestingly, when we maintain the exclusion of pairs where both groups of statistics agree (criterion 1), but we remove the exclusion of "matched" pairs (criterion 2), performance drops steeply to 58% (**Table S8**). Thus, even the "simpler" task of deciding between spectral and HOS when they explicitly disagree is difficult to achieve with good accuracy. Note that this low accuracy is despite the classes in this example being more balanced.

| | **Label: No HOS improvement** | **Label: HOS improvement** | |
|---|---|---|---|
| **Predicted: No HOS improvement** | 218 | 135 | 353 |
| **Predicted: HOS improvement** | 118 | 135 | 253 |
| | 336 | 270 | |

**Table S8.** Classification of pairs of texture image patches as being better segmented by HOS or not, where the pairs of patches with agreement between spectral statistics and HOS not included in training and testing, but both "matched" and "unmatched" pairs are included.

Also, when we remove criterion 1 (we include pairs where spectral and HOS agree), but we keep criterion 2 (we remove the "matched" pairs), we also observe a decrease in performance, although less pronounced than in the previous case, with accuracy falling to 77% (**Table S9**). This indicates that the major source of the low accuracy is in the mix of "matched" and "unmatched" pairs, and not because of the heterogeneity of pairs with no HOS improvement, or due to the dataset imbalance.

| | **Label: No HOS improvement** | **Label: HOS improvement** | |
|---|---|---|---|
| **Predicted: No HOS improvement** | 875 | 38 | 913 |
| **Predicted: HOS improvement** | 259 | 108 | 367 |
| | 1134 | 146 | |

**Table S9.** Classification of pairs of texture image patches as being better segmented by HOS or not, where "matched" pairs are excluded from training and testing, but pairs of patches with agreement between the two sets of statistics are included.

Thus, with these examples we show a control case where the method of analysis of fitting a model to the model outputs is expected to succeed, and we find a high accuracy for this case. We also show that the poor performance of the model is mostly due to the presence in the dataset of examples where HOS improve the segmentation task by favoring segmentation of pairs (unmatched pairs), and examples where they improve segmentation by disfavoring segmentation of pairs (matched pairs). We also show that excluding the pairs where spectral and HOS agree improves classification in the cases where we only train and test on the pairs with “unmatched”, but that this manipulation alone is not sufficient to reach high performance.

### S5) Low agreement between models of HOS fine-tuning and psychophysics:

One way to test the hypothesis that the visual system is fine-tuned to use HOS for segmentation in cases where they are more informative, is to identify or synthesize images where, from natural image statistics, we would predict HOS to be particularly useful for segmentation. Then, these could be used to probe human segmentation. Although such an empirical testing of this hypothesis is outside of the scope of this work, the results from our prior experimental work (Herrera-Esposito et al., 2021) allow for a preliminary exploratory analysis.

In this previous experimental work, despite observing an overall weak effect of HOS in segmentation, we observed some variation in the magnitude of the effect between the 4 different pairs of textures used. These were pairs of textures that had their spectral statistics matched, but that differed in their HOS. To test whether this variation between textures responds to a fine-tuning of the visual system to use HOS in some cases but not others, we analyzed whether the predictions from our segmentation models and from the models estimating usefulness of HOS for these pairs of textures relate to the observed variability in the psychophysical experiment.

For this, we first trained the linear model for segmentation on the BSD. Then, we tested the model on the pairs of textures used in our experimental work (Herrera-Esposito et al., 2021) that had matched spectral statistics but different HOS (experiment 3, Figure 6 in the original work), and extracted the linear activation from the model for each pair of textures. Because the models were trained with matched patches being the positive samples, we changed the sign of the linear output, so that higher (more positive) values indicate stronger segmentation. We repeated this procedure 20 times, because the 10-fold cross validation for choosing the scaling parameter (see above) adds some variability to the training procedure, and we report the mean and variation across models. The same procedure was also done using the models trained to identify pairs of patches with useful HOS. In this case, we did not have to change the sign of the linear output, since the positive samples were those with useful HOS. The same procedure for both kinds of models was also reproduced using the natural texture dataset instead of the BSD.

For the experimental data, we extracted the effect fitted to each texture using generalized linear mixed models, as described in the experimental study. Specifically, we extracted the interaction term between HOS dissimilarity and texture discontinuity from the full model, fitted to each texture individually. This fitted term is in units of log-odds ratio, and because of the coding of variables used in the statistical model, we also changed the sign so that larger (more positive) values indicated stronger segmentation (see original work for more detail).

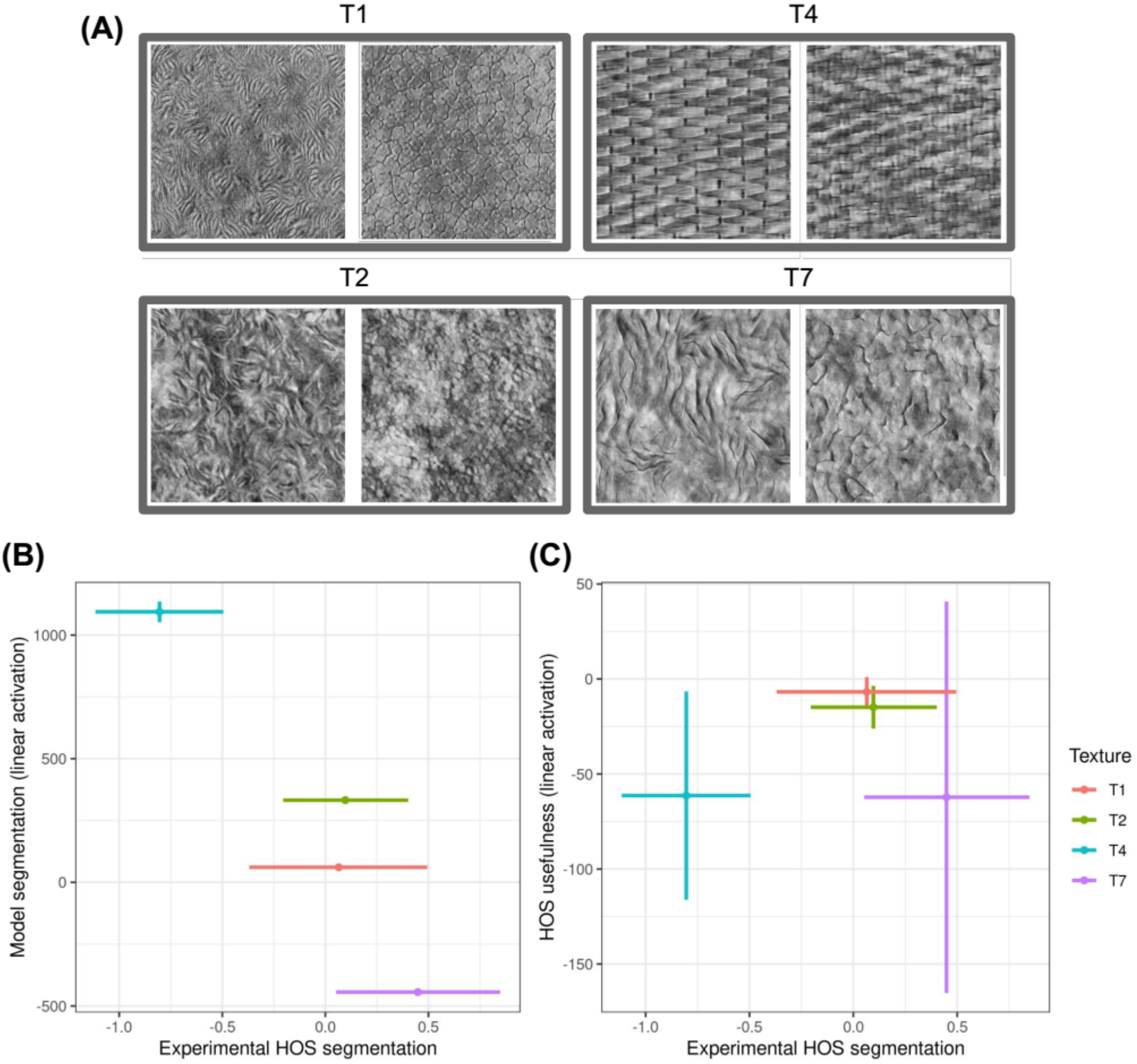


**Figure S2.** Comparison of experimental texture segmentation results (data obtained from (Herrera-Esposito et al., 2021)). The horizontal axis shows the estimated effect, on the segmentation of two patches of texture, of introducing a HOS mismatch between the two. Larger values indicate stronger segmentation when inducing the HOS mismatch. **(A)** Textures used in the experiment. Each gray rectangle groups together two patches of the textures that comprise a pair. **(B)** The vertical axis shows the linear activation component of the segmentation model using only HOS, trained on natural images and then tested on the textures of the experimental stimuli (see Methods). **(C)** The vertical axis shows the linear activation of the model trained to identify pairs of natural image patches where HOS are better than spectral statistics for segmentation. The texture numbering of the cited work is maintained in this figure, and all 4 textures used are shown.

In **Figure S2A** we show, for each pair of textures, the estimated empirical effect of HOS dissimilarity in human segmentation, and the linear component of the segmentation model using only the image HOS. We see that the strength of the segmentation shown by the model does not follow the observed experimental segmentation. This is the case also if we remove texture T4, which we argue to be an outlier in the original work, where we attribute the strong negative effect of HOS dissimilarity on segmentation to an artifact due to a phase effect (see further discussion in (Herrera-Esposito et al., 2021), Experiment 3). In **Figure S2B** we show the same analysis, using the linear component for the model predicting whether HOS perform better than spectral statistics (i.e. whether HOS should be used to segment the image). Again, we see that there is hardly any clear relation between experimental segmentation and model prediction. These results are similar when using models trained on textures, rather than natural images, and also when using both spectral and HOS together as predictors in the models.

Due to the small number of textures used in this analysis, the uncertainty in the estimates of the individual textures, and the fact that the experiment was not designed to test this hypothesis, this result should be taken with great caution. But the lack of a clear association between the observed human segmentation and the segmentation from models using HOS, or the estimate of HOS usefulness, may reflect that these HOS are a weak segmentation cue overall in peripheral vision.
